# A Principled Framework for Using Correlated Traits to Improve Risk Prediction

**DOI:** 10.64898/2026.08.23.746504

**Authors:** Kaiqian Zhang, Robert Bierman, Joshua M. Akey

**Affiliations:** Lewis-Sigler Institute for Integrative Genomics, Princeton University, Carl Icahn Laboratory, South Drive, Princeton, 08544, NJ, USA

**Keywords:** Polygenic score, Risk prediction, Correlated traits, Helper traits, Genetic correlation, Heritability, Type 2 diabetes, UK Biobank

## Abstract

**Background:** Although many complex phenotypes and diseases are influenced by shared genetic and environmental factors, risk prediction methods typically rely on genetic information from a single trait, leaving a rich source of predictive information largely unexploited. Phenotypic correlations can potentially be used to improve the accuracy of polygenic scores (PGS), but the conditions under which correlated traits meaningfully enhance prediction remain poorly understood. Here, we develop a general theoretical and simulation framework that quantifies the extent to which correlated “helper” traits improve predictive accuracy and identifies the factors that determine the magnitude of these gains.

**Results:** We show that helper traits can substantially improve predictive accuracy, with the magnitude of these gains governed by baseline model performance, genetic and environmental correlations, and the heritability of the target and helper traits, providing principled guidance for helper-trait selection. Paradoxically, when the target trait is itself weakly heritable, helper traits need not be highly heritable to substantially improve the accuracy of PGS, because low-heritability traits can still capture non-redundant environmental factors shared with the target trait. We empirically evaluated the use of helper traits by developing PGS models to predict type 2 diabetes using data from the UK Biobank. Helper traits substantially improved predictive accuracy relative to a single-trait PGS (AUC-ROC **= 0.907** versus **0.677**) and achieved performance comparable to models that use HbA1c (AUC-ROC **= 0.889**), the current clinical gold-standard biomarker.

**Conclusions:** Our results establish a general theoretical and practical framework for exploiting correlated traits to improve polygenic prediction, provide principled guidance for selecting informative helper traits, and demonstrate how shared genetic and environmental architecture can be leveraged to substantially increase predictive accuracy. Furthermore, we developed an interactive web application to estimate the expected gain in accuracy from candidate helper traits using empirically measurable quantities.

## Background

Polygenic scores (PGS) [1–3] use genetic variants identified through genome-wide association studies (GWAS) [4–6] to predict trait values and estimate disease risk. To date, most approaches for constructing PGS rely on information from a single trait [7–9]. However, a growing body of work has found pervasive correlations among complex phenotypes and diseases in humans [10–13], as well as widespread pleiotropy in GWAS data [10, 14–17]. For example, Bulik-Sullivan et al. [10] constructed an atlas of genetic correlations across hundreds of human diseases and traits using LD score regression [18], revealing extensive shared genetic architectures among seemingly distinct phenotypes. These observations suggest that correlated traits may provide an additional source of information for improving disease risk prediction. More practically, the use of correlated traits in predictive models is becoming increasingly feasible, as many large, deeply phenotyped cohorts, such as the UK Biobank (UKB) [19] and All of Us [20], have been developed.

Although PGS are typically constructed from a single trait, several studies have shown that incorporating additional correlated phenotypes or clinical variables can improve predictive accuracy across a range of diseases [21–26]. However, these analyses narrowly evaluated the gain in prediction accuracy in the context of specific target traits and therefore provide limited insight into the general factors that determine when and why a correlated trait is useful. A notable exception is the seminal work of Maier et al. [21], who showed that a multiple-trait predictor based on GWAS summary statistics can significantly improve predictive accuracy across a range of neuropsychiatric traits. Despite the important insights provided by Maier et al. [21], their framework focuses on leveraging genetic correlations estimated from GWAS summary statistics to improve predictive performance in specific trait settings. As such, it does not provide a general framework for understanding how genetic and environmental correlations, along with other variables, jointly determine when a correlated trait is informative for prediction.

To address this important gap in knowledge, we investigate the factors that determine whether correlated phenotypes, which we refer to as “helper traits”, meaningfully improve predictive accuracy. Although helper traits can formally enter a predictive model in the same manner as conventional covariates, they are conceptually distinct. Covariates are typically included because they are established risk factors or account for potential confounding or nuisance variation, whereas helper traits are correlated phenotypes selected specifically for their potential to add predictive information beyond what a baseline model captures. Our framework therefore focuses not simply on whether an additional variable is associated with the target phenotype, but on quantifying the complementary information it contributes and identifying the factors that determine the resulting gain in predictive accuracy.

Using both novel theory and comprehensive simulations, we show that the predictive accuracy of a target trait can be substantially improved by including one or more helper traits in the model. We also demonstrate that predictive gains from helper traits are governed by the interplay between multiple factors, including the performance of the baseline model, the magnitude of genetic and environmental correlations, and the heritabilities of helper and target traits. We also developed a redundancy-aware feature selection strategy to identify informative, minimally redundant helper traits, which we leveraged to construct models that predict type 2 diabetes in the UK Biobank [19]. We find that the inclusion of helper traits substantially improves predictive accuracy relative to single-trait PGS models and achieves performance comparable to, and in some cases exceeding, HbA1c [27], the current clinical gold-standard biomarker. Collectively, our results provide a general and principled framework for identifying and using correlated traits to improve risk prediction and investigating the shared genetic and environmental architecture underlying complex traits.

## Results

### Theoretical insights into when helper traits improve prediction

As illustrated in Figure 1A, correlations between traits arise from both shared genetic (*ρ_g_*) and environmental (*ρ_e_*) factors, which together determine the observed phenotypic correlation (*ρ_p_*). Large-scale biobanks [19, 20, 28–32] contain hundreds to thousands of phenotypes measured on the same individuals, creating widespread opportunities to leverage correlations between traits to improve prediction. A fundamental question is when and by how much correlated helper traits improve prediction beyond what is achieved with genetic predictors alone. To address this question, we developed a general theoretical framework that quantifies the expected increase in prediction accuracy obtained by incorporating helper traits into the prediction model. The framework considers a target trait (*Y_t_*), a baseline predictor (*B*; such as a PGS), and a helper trait (*Y_h_*). Figure 1B summarizes the prediction framework, in which a baseline PGS is augmented with one or more helper traits to improve prediction of the target phenotype.

**Fig. 1.**
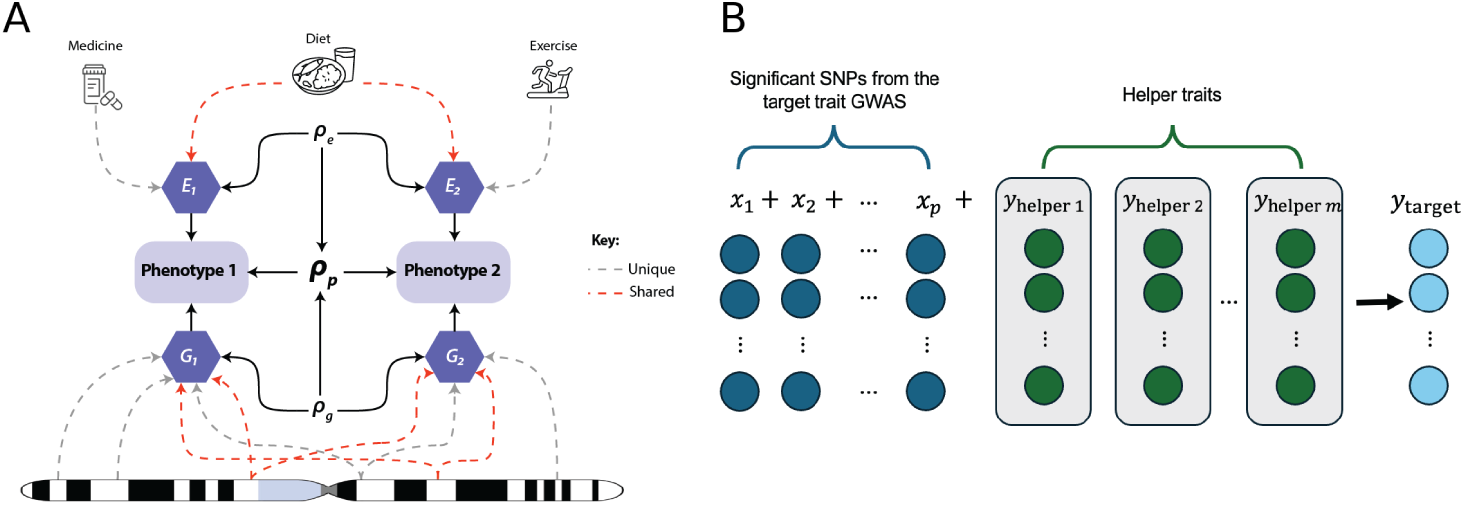
Schematic Representation of Leveraging Correlations Between Traits to Improve Predictive Accuracy. (**A**) Cartoon illustration showing how shared genetic (***G***) and environmental (***E***) factors for two traits contribute to phenotypic correlations. (**B**) Overview of a predictive model that leverages helper traits. SNPs significantly associated with the target phenotype (blue) and helper trait values (green) are jointly used to predict a target phenotype across individuals.

A central result of our theoretical analysis (derived in the Methods; Equation (13)) is that the relative gain in predictive accuracy 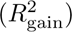 obtained by incorporating a helper trait into the model can be decomposed into three components: the total predictive signal provided by the helper trait, the redundancy of that signal with the baseline predictor, and the predictive performance of the baseline model:

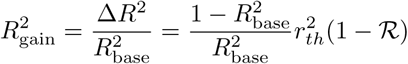

Here, 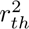 is the squared Pearson correlation between the helper trait (*Y_h_*) and the target trait (*Y_t_*) and represents the total predictive signal available from the helper trait. Likewise, 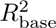 is the squared Pearson correlation between the baseline predictor (B) and the target trait, corresponding to the predictive accuracy of the baseline model. Thus, the factor 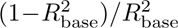 quantifies how much room there is to improve predictive accuracy and shows that helper traits provide the greatest benefit when the baseline predictor explains relatively little of the target phenotype. Finally, the redundancy parameter, R, quantifies the fraction of the helper trait’s predictive signal that is already captured by the baseline predictor, such that only the remaining non-redundant component contributes additional predictive information. An important implication of Equation (13) is that the marginal correlation between a helper and target trait is not sufficient to determine its predictive utility. Helper traits that have the same correlation with the target trait can differ substantially in the amount of non-redundant information they contribute beyond the baseline predictor, and therefore produce markedly different gains in predictive accuracy 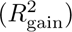.

To validate our theoretical results, we performed simulations across a broad range of parameter values and compared the observed gains in predictive accuracy with the theoretical expectations. In these simulations, each of the three determinants in Equation (13) was varied in turn while the other two were held fixed (Methods). As shown in Figure 2, the simulation results closely match the theoretical predictions across all scenarios examined. Consistent with Equation (13), predictive gain decreased as baseline predictive performance 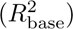 increased (Figure 2A), increased as the squared Pearson correlation between the helper and target trait 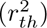 increased (Figure 2B), and decreased approximately linearly with the redundancy parameter (R) (Figure 2C). Across the parameter ranges examined, helper traits increased predictive accuracy by up to ∼ 70% relative to the baseline predictor, demonstrating that correlated traits can substantially improve prediction when they provide complementary information beyond what the baseline model already captures. Conversely, improvements became small when baseline predictive performance is high or when the information contained in the helper trait was redundant with the baseline predictor. Together, these results validate the theoretical decomposition of predictive gain and establish a general framework for understanding when correlated traits improve the prediction of polygenic phenotypes.

**Fig. 2.**
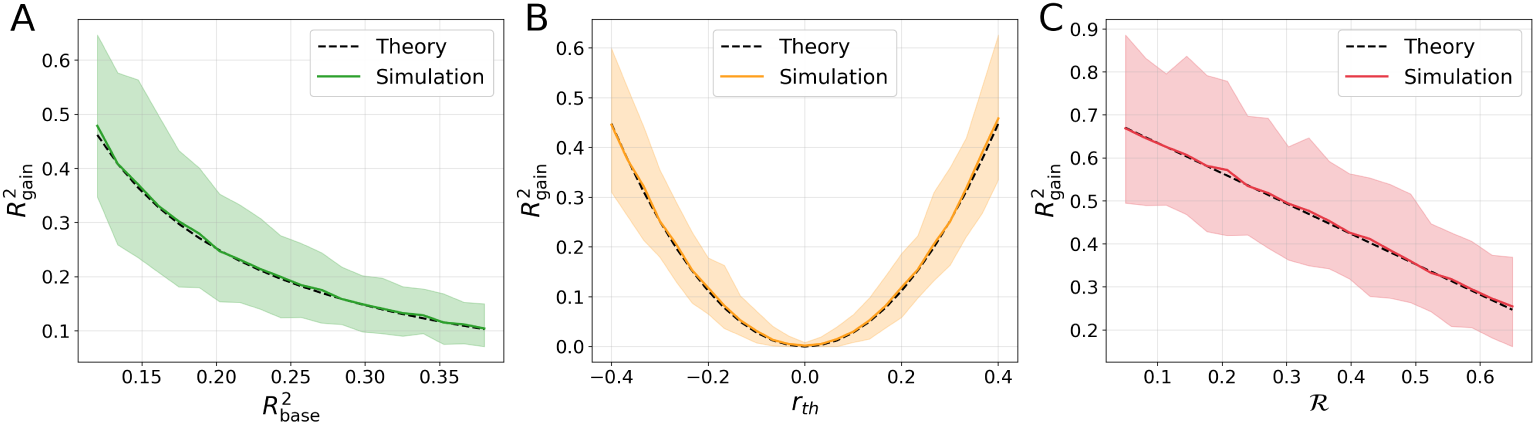
Simulations Validate Theoretical Results. The relative gain in prediction accuracy, 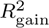, was evaluated in simulated data as a function of 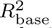 (**A**), *r_th_* (**B**), and R (**C**). Colored curves represent the simulated data and the shaded bands span the 2.5th to 97.5th percentile of the simulated values across replicates. Dashed black lines denote the corresponding theoretical predictions.

### Empirical insights into the characteristics of informative helper traits

Having established the theoretical determinants of predictive gain, we next investigated measurable characteristics that distinguish informative helper traits. Although Equation (13) identifies three factors that govern the expected gain in predictive accuracy of a helper trait —baseline predictive performance, correlation between helper and target traits, and redundancy—these quantities are themselves functions of the target and helper trait heritability and their genetic and environmental correlations. Equation (13) can be derived in terms of these measurable properties of helper and target traits (Additional file 1: Supplementary Methods 2), which we confirmed through simulations (Additional file 1: Figure S1). In the following, we also used the simulations to systematically evaluate how heritability and shared genetic and environmental factors influence gains in the predictive accuracy of helper traits.

We first investigated how the strength of shared genetic and environmental factors and helper trait heritability influences gains in predictive accuracy. In this scenario, we focus on a target trait that has high heritability (*h*^2^ = 0.9) and a baseline PGS model that only explains approximately 10% of target trait heritability (Figure 3). As expected, Figure 3A shows that when the target and helper traits share a strong genetic correlation (*ρ_g_* = 0.9), predictive gain increases with both the heritability of the helper trait and environmental correlation, with helper trait heritability exerting the stronger influence. This pattern is expected because the target trait is highly heritable, limiting the amount of predictive information that can be gained through shared environmental effects. Figure 3B presents the complementary analysis, in which the environmental correlation is fixed (*ρ_e_* = 0.9) and the genetic correlation is varied. Here the dependence is not monotonic. Because the helper–target phenotypic correlation is the sum of a genetic and an environmental contribution, 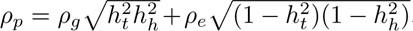, a negative genetic correlation opposes the fixed positive environmental correlation, and at the value of *ρ_g_* where the two contributions cancel the helper trait carries essentially no information about the target trait. This produces a valley of near-zero gain running through Figure 3B: at 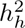 =0.9, for example, the gain falls from 6.41 at *ρ_g_* = −1 to essentially zero near *ρ_g_* = −0.1, then rises to 9.93 at *ρ_g_* = 1. Because a more heritable helper trait has a smaller environmental contribution to be cancelled, the valley shifts toward zero as helper heritability increases, from *ρ_g_* ≈ −1 at 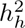 =0.1 to *ρ_g_* ≈ −0.1 at 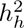 =0.9.

**Fig. 3.**
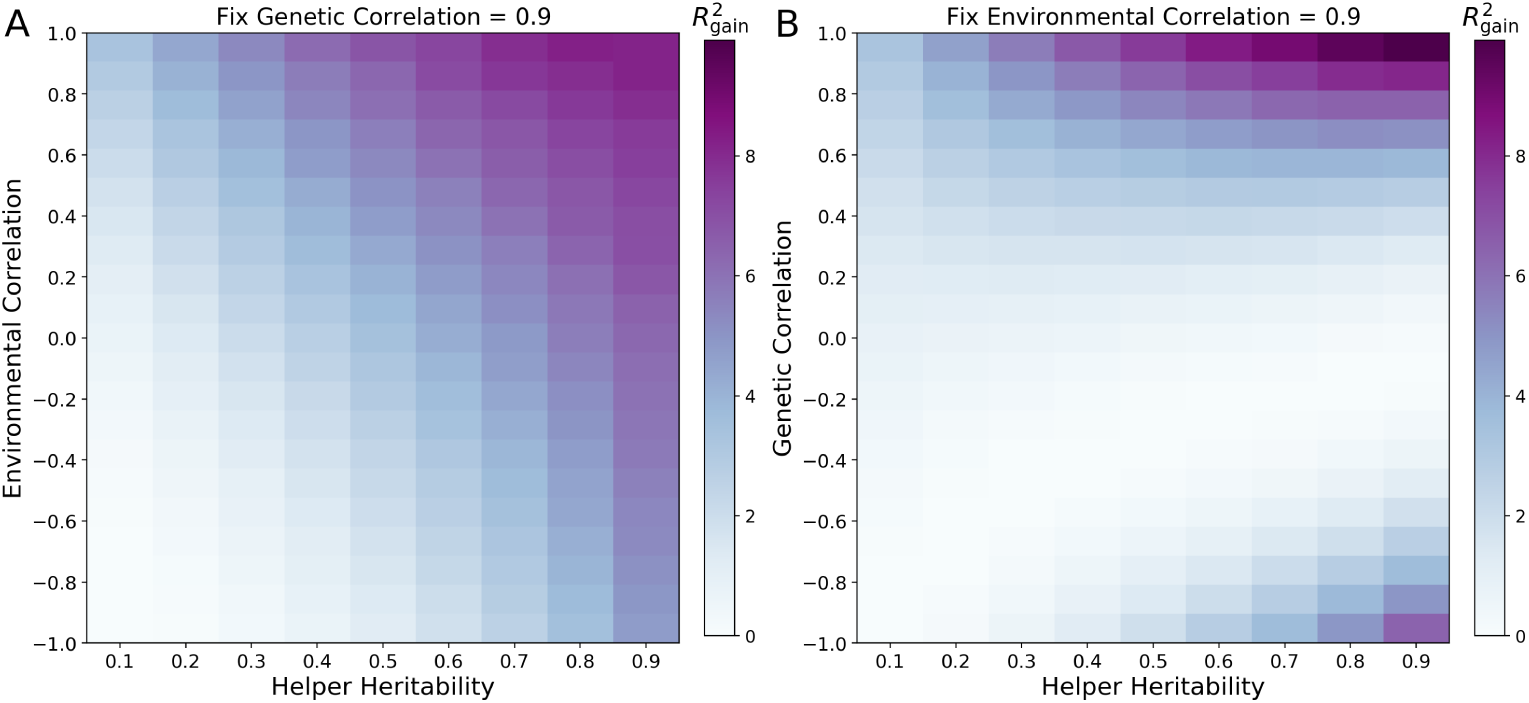
Gain in predictive accuracy for highly heritable target traits. The relative predictive gain 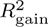for a target trait with 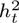 = 0.9 when the baseline predictor captures a fraction *α* = 0.1 of the target heritability. (A) The genetic correlation (*ρg*) is fixed at 0.9, and the environmental correlation (*ρe*) and helper heritability 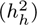 are varied. (B) The environmental correlation (*ρe*) is fixed at 0.9, and the genetic correlation (*ρ_g_*) and helper heritability 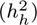 are varied.

The benefit of a more heritable helper trait therefore depends on where the trait sits in this landscape. Throughout Figure 3A, and away from the valley in Figure 3B, predictive gain increases with helper trait heritability, as expected from the theoretical framework (Additional file 1: Supplementary Methods 2), since a more heritable helper trait carries a larger share of genetic signal available to be shared with the target. Within the valley, however—for *ρ_g_* between approximately −0.3 and 0.3 in Figure 3B—increasing helper trait heritability moves the trait closer to exact cancellation, and the gain decreases. Both panels are also asymmetric about zero correlation, because correlations of the same sign reinforce one another whereas correlations of opposite sign partially cancel. That asymmetry is modest where the varied correlation contributes little to *ρ_p_* (in Figure 3A gains at *ρ_e_* = ±0.9 differ by roughly fivefold) and pronounced where it contributes most (in Figure 3B at 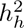 =0.5, *ρ_g_* = 0.3 gives a gain of 1.68 against 0.01 at *ρ_g_* = −0.3). In summary, when the target trait is highly heritable the largest gains come from helper traits that are themselves highly heritable and strongly positively correlated with the target phenotype. The exception is instructive: where a negative genetic correlation offsets a positive environmental one, the same increase in helper trait heritability instead reduces the gain. Even in this regime, therefore, the magnitude of the phenotypic correlation alone does not determine the value of a helper trait; how that correlation is composed also matters.

We next investigated whether the source of the correlation between the helper and target traits influences the gain in predictive accuracy when the target trait has low heritability. In contrast to Figure 3, which considered a highly heritable target trait (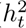 =0.9), Figure 4 examines a target trait with low heritability (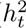 =0.1), where environmental factors account for most of its phenotypic variation. A natural expectation is that two distinct helper traits with the same phenotypic correlation to the target trait should produce similar improvements in predictive accuracy, regardless of how their correlations arise. Remarkably, our theory and simulations demonstrate that this intuition is incorrect. Specifically, Figure 4A shows that when the target and helper traits share a strong environmental correlation (*ρ_g_* fixed at 0.9, *ρ_e_*varied), helper traits with low heritability frequently produce the largest gains in predictive accuracy because they contribute information that is largely complementary to the baseline polygenic predictor. Figure 4B shows that this pattern persists when the environmental correlation is instead held at *ρ_e_*= 0.9 and the genetic correlation is varied: predictive gain declines with helper trait heritability at every value of *ρ_g_*, because the strong shared environmental component dominates the helper trait’s contribution. The dependence reverses only where the environmental correlation is weak. Along the rows of Figure 4A for which *ρ_e_* is close to zero, so that the helper-target correlation is almost entirely genetic, predictive gain increases with helper trait heritability, as it does throughout Figure 3A. The optimal helper trait heritability therefore depends on whether the helper-target correlation is predominantly environmental or predominantly genetic in origin, and not on its magnitude alone. These findings demonstrate that the benefit of a helper trait depends not only on the magnitude of the phenotypic correlation, but also on the genetic and environmental architecture underlying that correlation. Consequently, phenotypic correlation alone is not sufficient for identifying the most informative helper traits.

**Fig. 4.**
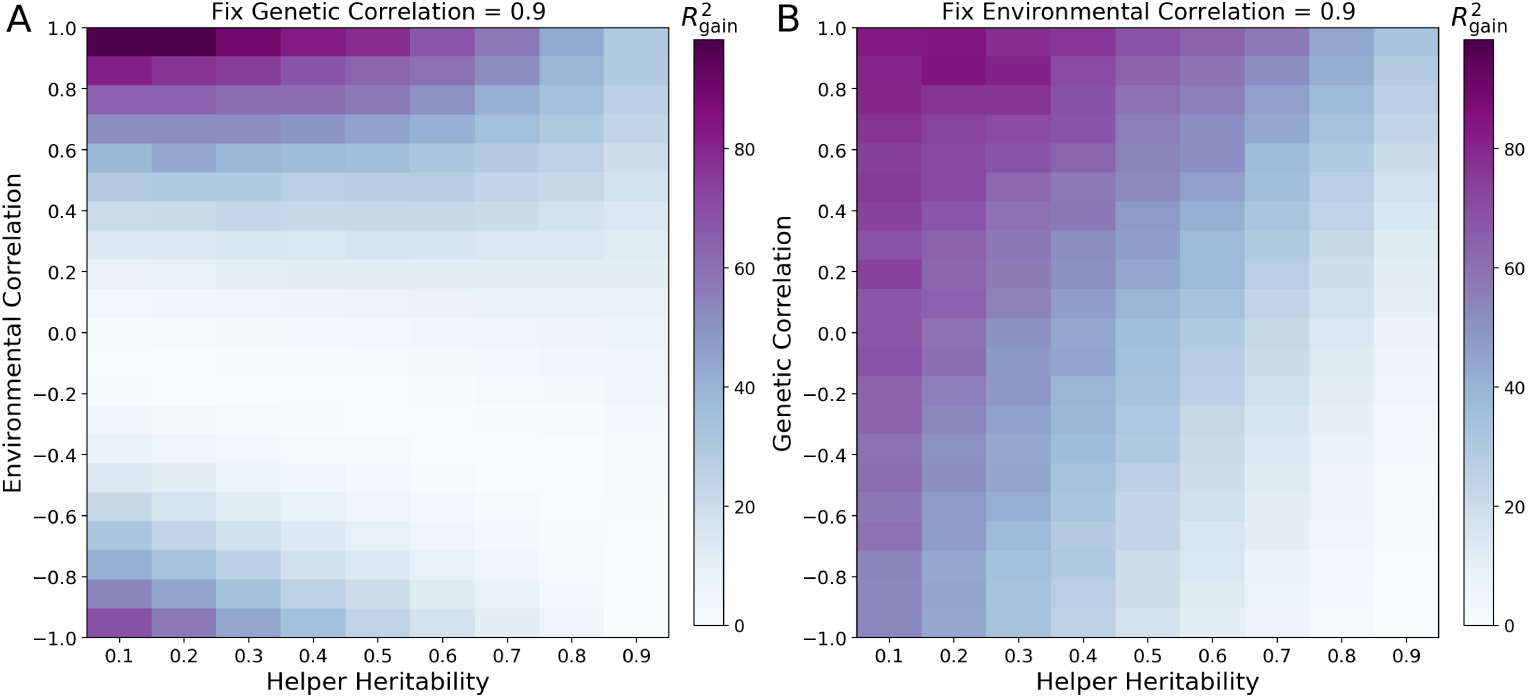
Gain in predictive accuracy for weakly heritable target traits. The relative predictive gain 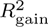 for a target trait with 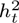 = 0.1 when the baseline predictor captures a fraction *α* = 0.1 of the target heritability. (A) The genetic correlation (*ρ_g_*) is fixed at 0.9, and the environmental correlation (*ρ_e_*) and helper heritability 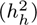 are varied. (B) The environmental correlation (*ρ_e_*) is fixed at 0.9, and the genetic correlation (*ρ_g_*) and helper trait heritability 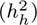 are varied.

An important practical question is whether helper traits remain useful once the baseline predictor is already highly accurate. Consistent with Equation (13), helper traits provided essentially no improvement when the target trait was highly heritable, and the baseline predictor already explained a large fraction of its variance (Additional file 1: Figure S2). In contrast, substantial gains in predictive accuracy remained possible in scenarios where the baseline predictor captures most of the target heritability (*α* = 0.9, so 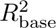 =0.09), but the target trait has low heritability (Additional file 1: Figure S3). In these cases, helper traits with low heritability and strong environmental correlation provide complementary information that is not available in the baseline polygenic predictor. These results further emphasize that the utility of helper traits depends critically on both the predictive performance of the baseline model and the characteristics of the helper and target traits.

### Application to type 2 diabetes: developing a baseline predictive model

To evaluate whether our theoretical framework improves disease prediction in empirical data, we applied it to type 2 diabetes (T2D) in the UK Biobank [19]. To develop a baseline predictive model, we first performed a genome-wide association study (GWAS) of T2D using REGENIE [33] in 11,411 cases and 132,718 controls and identified 37 independent genome-wide significant variants. These 37 SNPs and covariates, including sex, age, and the first 10 principal components (PCs) to account for population structure, were used to construct a baseline genetic predictor (see Methods). We then applied this baseline model to an independent dataset consisting of 81, 741 controls and 5, 876 cases, which was partitioned into three non-overlapping subsets: 20% to assess correlations among helper traits (see below), 60% for training, and 20% for testing. To fit the model, we used regularized logistic regression (Ridge) and obtained an AUC-ROC score of 0.677.

### Identifying and characterizing putative helper traits for type 2 diabetes

We assembled a comprehensive pool of 270 candidate helper traits consisting of 251 NMR-derived metabolomic traits [34] and 19 clinical and lifestyle traits previously associated with T2D [35–38]. This diverse collection of helper traits spans multiple biological processes implicated in diabetes, including glucose metabolism, lipid metabolism, inflammation, adiposity, cardiovascular physiology, and lifestyle-related risk factors. To better understand the relationship among putative helper traits, we first characterized their correlation structure (Figure 5A–C). For the 251 metabolomic traits, we constructed a network based on pairwise correlations (Figure 5A), which revealed numerous clusters corresponding to related biochemical pathways and lipoprotein subclasses. To explore heterogeneity in metabolite correlations, we quantified the number of strongly correlated neighbors (|*r*| *>* 0.8) for each NMR trait (Figure 5B). Traits ranged from having no highly correlated neighbors to over 40 (Figure 5B). Similarly, the 19 clinical and lifestyle traits formed biologically coherent clusters (Figure 5C), including strong associations among measures of adiposity, blood pressure, and lipid metabolism. Together, these data demonstrate that many candidate helper traits are highly redundant, suggesting that selecting traits solely on the basis of their individual association with T2D would introduce substantial overlap into predictive models.

**Fig. 5.**
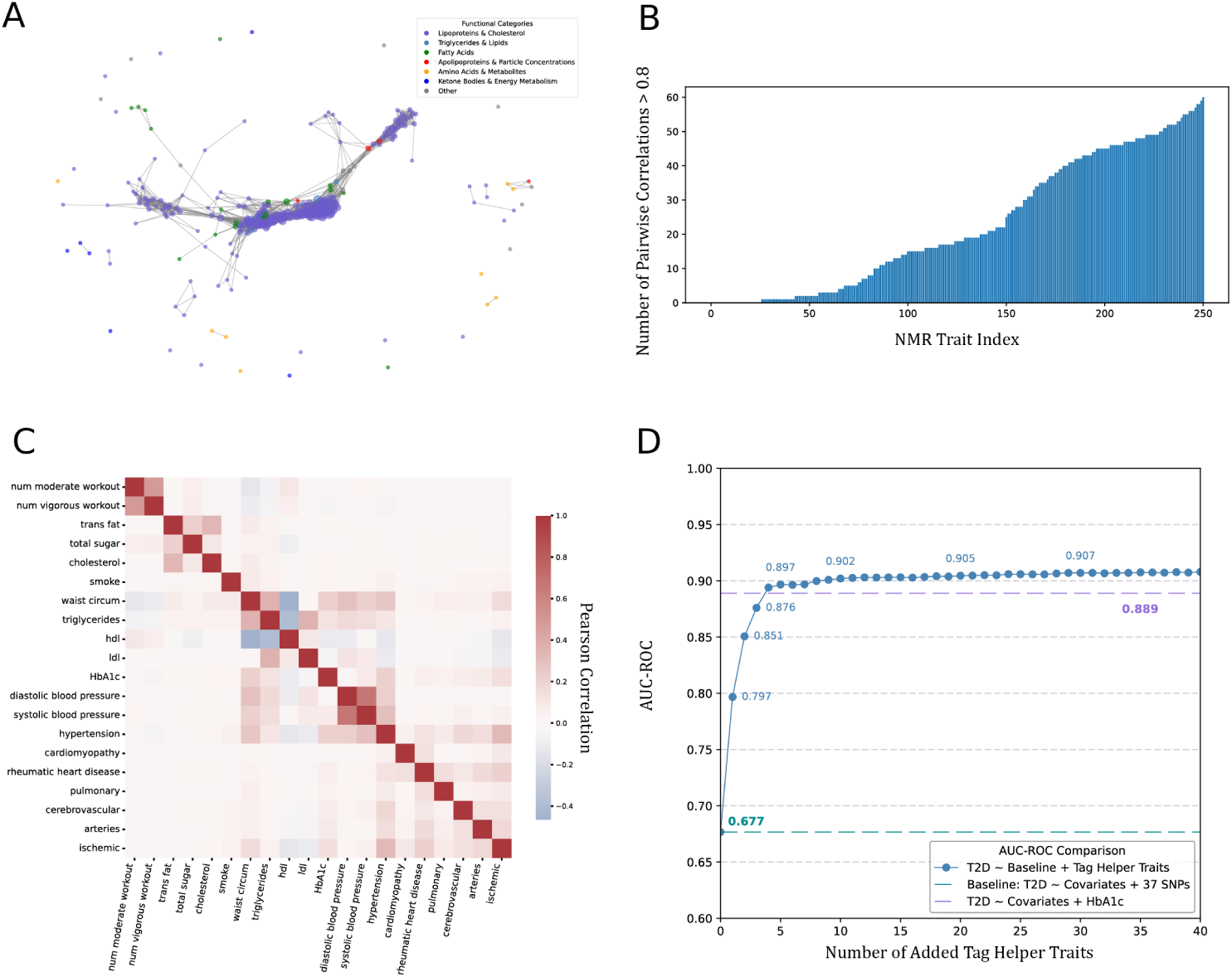
UK Biobank Results. (**A**) A network graph of NMR traits with strong correlations (|*r*| *>* 0.8), where nodes represent individual traits and edges indicate pairwise correlations above the threshold. Nodes are color-coded according to functional categories: lipoproteins & cholesterol (purple), triglycerides & lipids (blue), fatty acids (green), apolipoproteins & particle concentrations (red), amino acids & metabolites (orange), ketone bodies & energy metabolism (yellow), and others (gray). Highly interconnected clusters suggest biochemical redundancy among related traits, while sparsely connected nodes indicate unique metabolic signals.(**B**) The number of highly correlated neighbors (|*r*| *>* 0.8) for each NMR trait, providing a trait-wise view of correlation density. (**C**) is a heatmap of pairwise Pearson Correlation for 19 literature-based helper traits including lifestyle and clinical factors. (**D**) Each point on the x-axis indicates the number of top-ranked helper traits added to the baseline model (T2D ∼ Covariates + 37 SNPs), while the left y-axis shows the model’s AUC-ROC. The blue circles track performance as successive helper traits are introduced, quickly surpassing the baseline (green dashed line at AUC-ROC ≈ 0.677) and eventually exceeding the reference model using covariates plus HbA1c (brown dashed line at ≈ 0.889).

To efficiently identify a subset of helper traits that maximizes predictive information while minimizing redundancy, we developed a greedy algorithm to discover “tag helper traits” (Methods and Additional file 1: Supplementary Methods 1). The goal of this redundancy-aware algorithm is analogous to approaches designed to identify tag-SNPs for GWAS studies [39, 40]. Because our theoretical framework predicts that informative helper traits should exhibit strong correlations with the target phenotype, we first ranked candidate helper traits according to their correlation with HbA1c. HbA1c is a widely used clinical biomarker of glycemic control and T2D risk [27] and therefore provides greater statistical resolution than the binary disease phenotype for identifying informative helper traits. Applying this algorithm to the 270 candidate helper traits resulted in a set of 49 tag helper traits (Additional file 1: Figure S4) that were used in the analyses described below.

### Helper traits substantially improve the prediction of type 2 diabetes

To evaluate how the selected helper traits improve the accuracy of predicting T2D, we trained prediction models for T2D using AutoGluon [41], an automated machine learning framework that evaluates a diverse collection of linear and nonlinear prediction algorithms. As a baseline, we first fit a model containing the 37 significant variants identified by GWAS and covariates as described above. We then sequentially incorporated the 49 tag helper traits into the model in order of decreasing correlation with HbA1c, allowing us to quantify the incremental contribution of each helper trait to predictive performance (Figure 5D).

Incorporating helper traits into the model produced substantial improvements in predictive accuracy (Figure 5D). The baseline genetic model achieved an AUC-ROC of 0.677, whereas sequential addition of tag helper traits yielded rapid increases in predictive performance before gradually plateauing. Remarkably, incorporating only the single highest-ranked helper trait increased the AUC-ROC from 0.677 to 0.797 (Figure 5D). The rapid initial improvement in AUC-ROC followed by diminishing returns is consistent with our theoretical results. Specifically, the highest-ranked helper traits capture most of the available complementary predictive information, and subsequently added helper traits contribute progressively less because their information becomes increasingly redundant with what is already captured by the model. The final model with all 49 tag helper traits achieved an AUC-ROC of 0.907 (Figure 5D), substantially outperforming the baseline genetic model. These data show that a relatively small set of carefully selected helper traits can recover a large amount of predictive information beyond what is captured by PGS alone.

To place these improvements in a clinical context, we compared the helper-trait model to a model based on HbA1c (see Methods), the current gold-standard biomarker for T2D diagnosis and risk assessment [27]. The HbA1c model achieved an AUC-ROC of 0.889, indicating that the helper trait model slightly exceeded the predictive performance of this established clinical biomarker (Figure 5D). Thus, although many complex diseases lack a highly informative biomarker such as HbA1c for T2D, our results suggest that a carefully constructed set of helper traits provides a practical strategy for developing predictive models whose performance is commensurate with those based on established clinical biomarkers.

## Discussion

Predicting complex traits and disease risk has traditionally focused on extracting increasingly sophisticated information from genetic variation identified by GWAS [3, 42, 43]. Here, we demonstrate that substantial gains in predictive accuracy can be achieved by systematically leveraging phenotypically correlated helper traits. Although the intuitive appeal of incorporating correlated phenotypes has long been recognized [21, 44–46], the conditions under which helper traits improve prediction—and how they should be selected—have remained poorly understood. Our study addresses this gap by developing a theoretical framework that identifies the key factors that make helper traits informative and by translating these insights into a practical strategy for improving disease prediction.

Helper traits should be conceptually distinguished from conventional covariates in predictive models. Although both can be incorporated mathematically as additional predictors, helper traits are explicitly selected based on their relationship with the target phenotype and their potential to contribute information not already captured by the baseline predictor. Our theoretical framework quantifies this additional information and therefore provides a principled basis for determining when adding a correlated phenotype meaningfully improves prediction, rather than treating all available covariates as interchangeable candidate predictors.

A central finding of our study is that the benefits of a helper trait are governed not simply by its phenotypic correlation with the target trait, but by how that correlation arises. Our theoretical framework demonstrates that predictive gain depends on three quantities: baseline predictive performance, correlation between the helper and target phenotypes, and redundancy between the helper trait and the baseline predictor. We further showed that these quantities are themselves determined by biological properties (Additional file 1: Supplementary Methods 2), including the heritability of the target and helper traits and their underlying genetic and environmental correlations. Notably, these properties can be estimated with existing methodological tools, including SNP-based variance-component methods and LD-score regression for estimating heritability and genetic correlation, and multivariate twin or family models for decomposing phenotypic covariance into genetic and environmental components [10, 47–49]. Collectively, these results provide a conceptual framework for understanding why some correlated traits substantially improve prediction whereas others contribute little additional information.

We selected T2D as an empirical application because it provides an ideal setting for evaluating helper trait based prediction. T2D is a major public health challenge worldwide [50, 51] and has been extensively characterized through large-scale genetic and epidemiological studies [52–54], enabling both well-powered GWAS for constructing baseline polygenic predictors and the identification of a rich collection of candidate helper traits spanning metabolomic, clinical, and lifestyle domains [19, 35, 55]. Applying the theoretically derived principles to T2D in the UK Biobank further demonstrated the practical benefit of helper traits. Specifically, we found that many candidate helper traits contained highly redundant information, motivating the development of a redundancy-aware feature-selection algorithm (Additional file 1: Supplementary Methods 1) to identify sets of minimally redundant “tag helper traits.” Incorporating the selected helper traits substantially improved predictive performance, increasing the AUC-ROC from 0.677 for the baseline genetic model to 0.907. Our findings are consistent with those of He et al. [56], who constructed a polyexposure score from non-genetic exposures for T2D and found that it discriminated substantially better than a polygenic score (C statistic 0.762 versus 0.709), although it added only modest incremental value beyond established clinical risk factors (C statistic 0.839). More broadly, these findings illustrate that carefully selected helper traits can recover substantial predictive information that is not captured by PGS, highlighting the value of integrating genetic and phenotypic information within a unified prediction framework [24, 46].

An intriguing question is whether helper traits can improve the transferability of PGS across populations [57]. Indeed, one of the major challenges facing polygenic prediction is the marked decline in predictive accuracy observed when models are applied across genetically diverse populations [58–61]. Because helper traits incorporate phenotypic information beyond genetic variation alone, they may capture complementary signals that are more robust across populations than polygenic risk scores themselves. Determining whether theory-guided selection of helper traits can mitigate ancestry-related declines in predictive performance represents an important direction for future investigation.

Our study has several limitations. First, the theoretical framework considers pairwise relationships between a target trait and a single helper trait, whereas real-world prediction often involves simultaneously integrating many correlated helper traits. Although we addressed this challenge empirically through redundancy-aware feature selection, extending the theoretical framework to explicitly model multiple helper traits remains an important direction for future work. Second, our empirical analyses were performed using UK Biobank, and the optimal set of helper traits will likely vary across populations and datasets with different phenotypic measurements. Finally, although our predictive models were optimized using automated machine learning, future work should investigate how interpretable modeling approaches can better elucidate the biological mechanisms underlying the observed improvements in predictive accuracy.

## Conclusions

Large-scale biobanks are increasingly collecting thousands of quantitative phenotypes spanning clinical measurements [19, 62], metabolomics profiles [63], proteomics measurements [64], imaging-derived phenotypes [65], wearable sensors data [66], and longitudinal electronic health records [67]. These rich phenotypic resources represent an underutilized source of complementary predictive information. By providing both a theoretical understanding of the potential power of helper traits and a practical framework to select them, our study establishes a general strategy for integrating correlated phenotypes with PGS to improve the prediction of complex and quantitative traits and diseases.

## Methods

### Statistical model and notation

To understand how correlated helper traits improve prediction accuracy, we developed a tractable statistical framework to describe the joint distribution of two correlated phenotypes. Let ***Y*** *_t_* and ***Y*** *_h_* denote the target and helper trait, respectively. We assume both traits follow Fisher’s polygenic model [68] with additive genetic effects and are standardized to have Normal distributions with means E[***Y*** *_t_*] = E[***Y*** *_h_*] = 0 and variances Var(***Y*** *_t_*) = Var(***Y*** *_h_*) = 1. Let ***X*** ∈ R*^n^*^×*p*^ denote the genotype matrix for *n* individuals and *p* genetic variants, with *X_ij_* ∈ {0, 1, 2} representing the genotype of individual *i* at variant *j*. We assume *X_ij_* ∼ Binomial(2*, p_j_*), where *p_j_* is the allele frequency of the *j*th variant. Let ***β****_t_* and ***β****_h_* be vectors of causal variant effect sizes for the target and helper traits, respectively, and let ***ε****_t_* and ***ε****_h_* denote the environmental noise terms for the target and helper traits, respectively. The target and helper traits can therefore be written as the sum of genetic (***G*** = ***Xβ***) and environmental (***E*** = ***ε***) components:

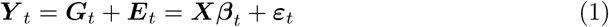

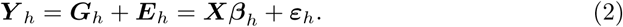

In the following, we use *ρ* to denote population-level correlation parameters, *r* to denote correlations estimated from observed variables, and *R*^2^ to denote the coefficient of determination from linear models.

### Decomposing phenotypic variation into its component sources

From Fisher’s additive polygenic model, the joint distribution of the target and helper traits is fully determined by the variances and covariances of the genetic and environmental effects. Therefore, we briefly show how phenotypic variation can be decomposed into its component sources and derive an expression for the phenotypic correlation between the target and helper trait that will facilitate simulations. We assume that genetic and environmental effects are independent, i.e., Cov(***Xβ****, **ε***) = 0, so that the covariance between traits can be written as the sum of genetic and environmental contributions: Cov(***Y*** *_t_, **Y** _h_*) = Cov(***Xβ****_t_, **Xβ**_h_*)+Cov(***ε****_t_, **ε**_h_*). We parameterize these quantities in terms of heritability and correlations. Specifically, we define the heritability of the target and helper traits as 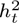 = Var(***Xβ****_t_*) and 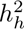 = Var(***Xβ****_h_*), respectively, so that Var(***ε****_t_*) = 1 − 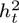 and Var(***ε****_h_*) = 1 − 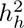 [69].

The genetic and environmental correlations, denoted as *ρ_g_*and *ρ_e_*, respectively, can be shown to be:

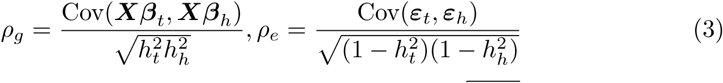

Rearranging these expressions yields Cov(***Xβ****_t_,* ***Xβ****_h_*) = 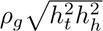 and Cov(***ε****_t_,* ***ε****_h_*) = 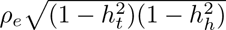 Using these definitions, the joint distributions of the genetic and environmental components can be written as:

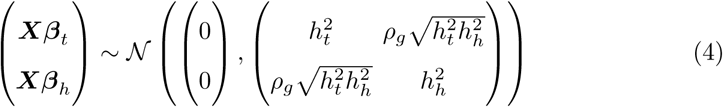

and

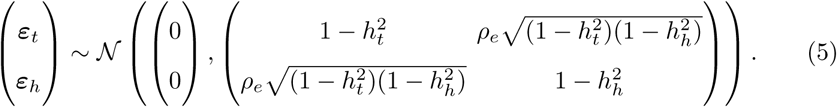

Finally, because ***Y*** *_t_*and ***Y*** *_h_*are sums of independent genetic and environmental components, their joint distribution is also multivariate normal with covariance given by the sum of these two matrices. This yields:

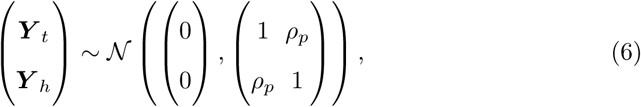

where the off-diagonal element *ρ_p_* is the phenotypic correlation between the target and helper traits. Because *ρ_p_* is the sum of the genetic and environmental contributions, the theory described above allows us to derive a key relationship linking phenotypic, genetic, and environmental correlations with the heritability of the target and helper traits:

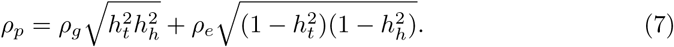

In words, Equation (7) shows that the phenotypic correlation between two traits is a weighted sum of their genetic and environmental correlations, with the weights determined by functions of the heritability’s of the two traits. It also demonstrates that traits with similar phenotypic correlations may differ substantially in the extent to which their correlation is driven by shared genetic versus environmental factors. Thus, *ρ_p_* alone does not determine whether a helper trait provides information beyond what is captured by the baseline (single-trait) genetic predictor. Although we do not explore this point here, it is also worth noting that Equation (7) provides a way to estimate the environmental correlation between traits, assuming unbiased estimates of *ρ_p_, ρ_g_,* 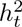 and 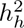 can be obtained. Thus, the theory described above may also be useful in developing approaches to better understand shared non-genetic determinants of disease.

### Expected gain in prediction accuracy

We derived an expression to quantify the improvement in prediction accuracy from a helper trait relative to a baseline model without it. Let ***B*** denote a standardized baseline predictor model that includes only genetic data for the target trait. Specifically, the baseline model corresponds to a simple PGS [7], which constructs a predictor for each individual as a simple linear combination of variants significantly associated with the target trait, weighted by their effect sizes: *B_i_* = ∑*_j_ ^β_j_ x_ij_*, where *^β_j_* and *x_ij_* are the estimated effect size and genotype of individual *i* at variant *j*. Let *r_th_* = Cor(***Y*** *_t_,* ***Y*** *_h_*)*, r_tB_* = Cor(***Y*** *_t_,* ***B***), and *r_hB_* = Cor(***Y*** *_h_,* ***B***) denote the Pearson correlation coefficients between the target trait and helper trait, target trait and baseline predictor, and helper trait and baseline predictor, respectively.

To quantify the gain in predictive accuracy from incorporating the helper trait, we consider the joint linear model ***Y*** *_t_* = *β*_0_ + *β_B_**B*** + *β_h_**Y** _h_* + ***η***, where *β*_0_ is the intercept term (which equals zero when variables are standardized), *β_B_* and *β_h_* are regression coefficients that quantify the contributions of the baseline predictor and helper trait to the prediction of the target trait, and ***η*** denotes the residual term orthogonal to both ***B*** and ***Y*** *_h_*. Note that the baseline predictor ***B*** represents a PGS constructed from multiple variants and is treated as a single scalar predictor in the regression model. Because all variables are standardized, the coefficient of determination, *R*^2^, for the joint model is the squared multiple correlation between ***Y*** *_t_* and the linear predictor defined by (***B****, **Y** _h_*), which can be written as:

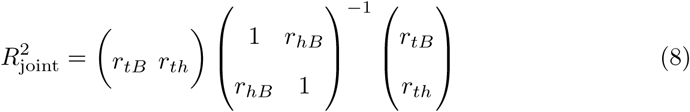

and evaluating the inverse yields:

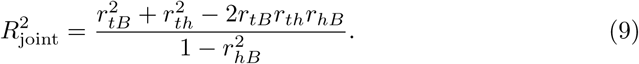

The absolute improvement in predictive accuracy from adding the helper trait is therefore:

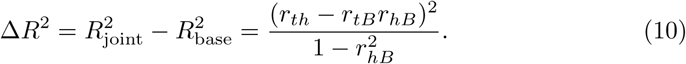

where 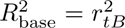, because the coefficient of determination for a single standardized predictor is equal to the squared Pearson correlation. Although Equation (10) provides an exact expression for the gain in predictive accuracy, its nonlinear dependence on multiple pairwise correlations obscures its interpretation. In particular, the numerator *r_th_* − *r_tB_r_hB_* reflects the portion of the helper-trait association that is not explained by the baseline predictor, but this quantity is not scale-invariant and depends on the marginal variances explained by both ***B*** and ***Y*** *_h_*.

To obtain a more interpretable quantity, we consider the correlation between ***Y*** *_t_*and ***Y*** *_h_* after accounting for their shared dependence on ***B***. This is precisely what is captured by the partial correlation between the target and the helper trait conditional on the baseline predictor:

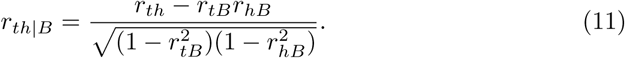

The partial correlation, therefore, provides a normalized measure of the non-redundant association between the helper and target traits after accounting for the baseline predictor. After squaring equation (11) and substituting it into equation (10), we obtain:

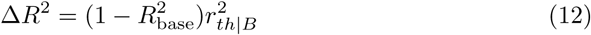

Equation (12) shows that the gain in predictive accuracy depends on the squared partial correlation between the helper and target trait. However, 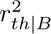 conflates two distinct factors: the overall strength of association between the helper and target traits and the extent to which this association is already captured by the baseline predictor. In particular, two helper traits with identical marginal correlations *r_th_* may yield different predictive gains depending on how much of their signal is redundant with ***B***. To disentangle these effects, we reparametrize the partial correlation in terms of a redundancy measure that quantifies the fraction of the helper-target association explained by the baseline predictor. Specifically, we define 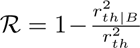, where R ≤ 1. R quantifies the fraction of the marginal correlation between the helper and target trait that is redundant with the baseline predictor. When R = 0, the helper trait provides entirely independent information about the target trait, whereas R = 1 indicates that all of the helper-target association is redundant with the baseline predictor. Negative values of R are also possible, and arise when conditioning on ***B*** strengthens rather than weakens the helper-target association, a classical suppression effect. In that regime the helper trait contributes more alongside the baseline predictor than its marginal correlation alone would suggest. Substituting the redundancy equation into equation (12) yields 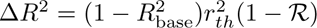. Finally, we define the relative gain in predictive accuracy as:

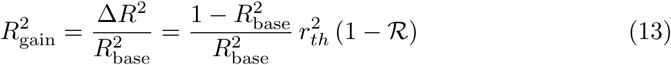

Equation (13) provides a simple interpretation of the relative predictive benefit of a helper trait. The gain increases with (i) weaker baseline predictive performance through the factor 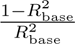, (ii) stronger marginal correlation between the helper and target trait through 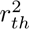, and (iii) lower redundancy with the baseline predictor, through (1 − R). This parameterization makes it clear that helper traits with similar marginal correlations to the target trait may differ substantially in utility, depending on how much of their signal is redundant with the baseline predictor. In Additional file 1: Supplementary Methods 2, we derive an expression for Equation (13) that parameterizes 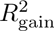 in terms of the target and helper trait heritabilities and their genetic and environmental correlations. To facilitate use of our theoretical framework by the broader scientific community, we developed an interactive web application (https://akeylab.github.io/correlated-traits-prediction/) that enables investigators to estimate the expected gain in predictive accuracy for candidate helper traits using empirically measurable quantities, including target- and helper-trait heritabilities, genetic and environmental correlations, and baseline predictive performance.

### Simulations

#### Specifying trait architecture and correlation structure

We developed a framework to simulate a target trait ***Y*** *_t_*, helper trait ***Y*** *_h_*, and baseline polygenic predictor, ***B***, from the following five parameters: the heritability of the target and helper traits (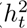 and 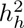), the genetic and environmental correlation between ***Y*** *_t_* and ***Y*** *_h_* (*ρ_g_* and *ρ_e_*), and the expected baseline predictive performance 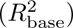 . The phenotypic correlation, *ρ_p_*, is determined by 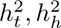, *ρ_g_,* and *ρ_e_* through Equation (7).

We simulated target and helper trait values according to the model defined in Equations (1) and (2). Thus, we need to specify effect sizes for causal variants that contribute to heritable variation in the target and helper traits. Unless noted otherwise, we assumed that 100 causal variants contributed to both the target and helper traits, approximating a polygenic architecture while keeping simulations computationally tractable. The helper trait was also influenced by an additional 10 helper trait-specific causal variants, providing flexibility in tuning the amount of helper trait signal that is independent of the target trait. The effect-size vectors ***β****_t_* and ***β****_h_* were constructed from three orthonormal basis vectors ***a***, ***b***, and ***c***, embedded in the same 110-dimensional space of causal variants (the 100 shared variants plus the 10 helper trait-specific variants). These basis vectors are not themselves the final effect sizes; rather, they define independent components of genetic effects that are scaled and combined to produce ***β****_t_* and ***β****_h_*. The basis vectors ***a*** and ***b*** were defined only for the shared causal variants by setting their entries for the 10 helper trait specific variants to zero, whereas ***c*** was defined for only the helper trait specific variants by setting its entries for the 100 shared variants to zero. The vectors ***a*** and ***b*** were constructed to be orthogonal. This decomposition separates helper-trait genetic effects into a component shared with the target trait (***a***) and two independent helper-specific components (***b*** and ***c***). Consequently, only ***a*** contributes to the genetic covariance between the target and helper traits, whereas ***b*** and ***c*** contribute only to helper-trait genetic variance. The effect-size vectors for the target and helper traits are calculated as:

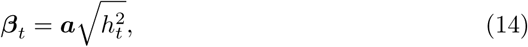

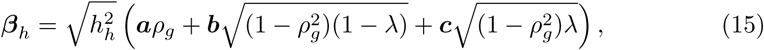

where *λ* ∈ (0, 1) controls the proportion of helper trait variance attributable to helper trait-specific variants. Unless otherwise noted, we set *λ* = 0.20. When genotypes are standardized, this construction yields Var(***Xβ****_t_*) = 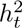, Var(***Xβ****_h_*) = 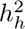, and Cor(***Xβ****_t_, **Xβ**_h_*) = *ρ_g_*.

Genotypes were simulated under a standard allele-count model. For each variant *j*, we sampled a minor allele frequency *p_j_* ∼ Uniform(0.05, 0.5), and generated individual genotypes as *X_ij_* ∼ Binomial(2*, p_j_*), where *X_ij_* ∈ {0, 1, 2} denotes the genotype of individual *i* at variant *j*. Genotypes were standardized to have mean zero and unit variance, *G_ij_* = (*X_ij_*− 2*p_j_*)*/*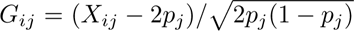 . Environmental residuals were then generated from a bivariate normal distribution:

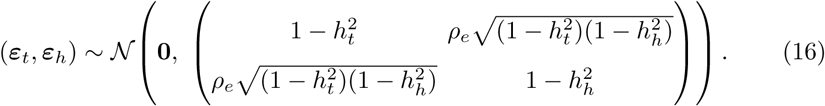

Phenotypes for each individual were then formed by adding the genetic and environmental components, ***Y*** *_t_* = ***G****_t_* + ***ε****_t_* and ***Y*** *_h_* = ***G****_h_* + ***ε****_h_*, and standardized to have mean zero and unit variance within each simulated dataset. The empirical correlation between the target and helper trait within each simulated dataset is denoted by *r_th_*= Cor(***Y*** *_t_, **Y** _h_*).

Finally, we derived an expression for the baseline predictor (or baseline PGS), ***B***, that captures a specified fraction of the target trait heritability. Specifically, we define 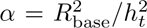, where *α* denotes the fraction of the target trait heritability captured by the baseline predictor. This framework allows us to recapitulate the fact that PGS developed from GWAS account for only a fraction of the target trait heritability (because GWAS does not typically discover all causal variants). The baseline predictor is then calculated as:

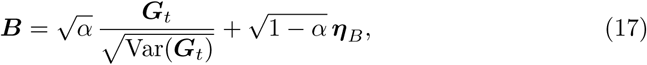

where ***η****_B_* is a random variable sampled from a N(0, 1) distribution. This construction expresses the baseline predictor as a weighted combination of the standardized genetic component of the target trait and Gaussian noise. Because both the standardized genetic signal and the independent noise have unit variance, the coefficients are chosen as √*α* and √1 − *α* (rather than *α* and 1 − *α*) so that the genetic signal contributes an *α* fraction of the variance of the baseline predictor, while the independent noise contributes the remaining 1−*α* fraction. Consequently, the baseline polygenic score has unit variance and an expected predictive performance of 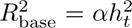. Note that, by construction, 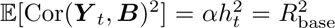, which is feasible only when 0 ≤ *α* ≤ 1, or equivalently, when 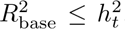. Parameter combinations violating this constraint were excluded.

In each simulation replicate, we calculate the empirical gain in predictive accuracy as:

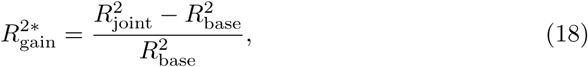

where 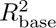and 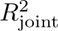 are the coefficients of determination obtained by fitting the baseline regression model ***Y*** *_t_* ∼ ***B*** and the joint regression model ***Y*** *_t_* ∼ ***B*** + ***Y*** *_h_*, respectively. We denote the empirical gain as 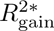 to emphasize that it is calculated in simulated data, whereas Equation (13) describes the theoretical equivalent metric 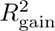.

#### Simulation to validate theoretical predictions

To validate the decomposition in Equation (13), we varied each of its three determinants in turn while holding the other two fixed: 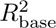 over [0.12, 0.38] at 20 evenly spaced values (with *r_th_* = 0.30 and R = 0.30), *r_th_* over [−0.40, 0.40] at 25 values (with 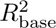 = 0.20 and R = 0.30), and R over [0.05, 0.65] at 20 values (with 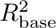 = 0.20 and *r_th_* = 0.42). Because Equation (13) is expressed entirely in terms of 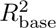, *r_th_* and R, this validation does not require simulated genotypes: for each parameter combination we drew the target trait, helper trait and baseline predictor directly from a standardized trivariate Normal distribution whose pairwise correlations were set to *r_tB_* = 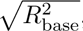, the specified *r_th_*, and the value of *r_hB_* that yields the requested redundancy R. Each parameter combination used 150 replicates of 2,000 individuals. Within each replicate, 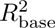 and 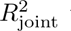 were computed in sample from the realized correlations and the empirical gain 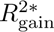 obtained from Equation (18); for two standardized predictors this squared multiple correlation has a closed form, so it was evaluated directly rather than by numerically fitting the two regressions. The theoretical gain 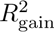 was calculated from Equation (13). The genotype-based framework described above is instead used for the analyses in Figures 3 and 4, and to validate the closed-form expression in Additional file 1: Figure S1.

#### Simulations to evaluate the effect of helper trait characteristics on predictive accuracy

To understand how characteristics of the helper trait influence prediction accuracy, we performed simulations across a broad range of genetic and environmental parameter values. Specifically, we varied 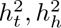 *, ρ_g_,* and *ρ_e_* while fixing *α*, the fraction of target heritability captured by the baseline predictor (so that 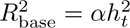). For each parameter combination, we simulated 20, 000 individuals, fit the baseline and joint regression models in-sample (without a train/test split), and computed the empirical relative gain in predictive accuracy, 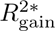, from the corresponding 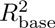 and 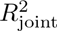 values using Equation (18).

#### Identifying ”tag helper traits”

We developed a greedy algorithm (Additional file 1: Supplementary Methods 1) to identify a set of “tag helper traits” that are minimally redundant with one another. We first calculate the absolute Pearson correlation between each candidate helper trait in the set *U* and the target trait, and rank the candidates in descending order of that correlation. Starting with the most strongly correlated helper trait, we iteratively add it to the selected set if its absolute pairwise correlation with all previously selected traits is below a predefined redundancy threshold *τ* . We used *τ* = 0.6 and did not cap the number of selected traits, so selection continued until no remaining candidate satisfied the threshold. This procedure yields a set of helper traits that are both strongly associated with the target trait and minimally redundant with one another. Because helper traits are ranked by decreasing correlation with the target trait, the algorithm preferentially retains the most informative representative from each cluster of correlated traits.

#### Type 2 diabetes data and GWAS

We applied the helper trait framework to predict type 2 diabetes (T2D) among individuals in the UK Biobank [19] (Field ID: 41270). Access was provided under application number 104628. We first performed GWAS for T2D using REGENIE [33]. After QC and filtering as previously described [70], we retained 11, 411 cases and 132, 718 controls. Imputed genetic data for these individuals were obtained from UKB for autosomal chromosomes (Field ID: 22828) and filtered with PLINK2 [71] for standard genotyping QC criteria, as previously described [70]. In total, 908 SNPs in the imputed dataset reached the genome-wide significance threshold of *p <* 5 × 10^−8^. We then performed linkage disequilibrium clumping at a threshold of 0.1 to identify independent associations. In total, we identified 37 independent SNPs significantly associated with T2D.

#### Helper traits for type 2 diabetes

We identified helper traits for T2D from two primary sources. First, because T2D is a metabolic disorder that produces widespread changes in circulating metabolites [55, 72], metabolomic traits provide a rich source of information about disease status. Accordingly, we analyzed 251 metabolomic traits (Field ID: 23475) measured in UK Biobank individuals using NMR profiling [34]. Second, we identified 19 clinical and lifestyle traits from a literature review [35–38] that were available among UKB participants. Because many of these traits are highly correlated with one another, naively including all of them in a predictive model would introduce substantial redundancy, leading to diminished performance and potentially unstable model estimates. We therefore used the algorithm described above to identify tag helper traits.

#### Predicting type 2 diabetes and comparing predictive models

The GWAS results were used to construct the baseline model: T2D ∼ Sex + Age + 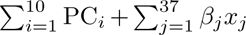, where T2D status was coded as 0 for controls and 1 for cases. The model includes sex, age, and the first 10 principal components (PCs) provided by the UKB (Field ID: 26201) to account for population structure, and a PGS defined as a weighted sum of the 37 variants identified by GWAS, where *x_j_* ∈ {0, 1, 2} denotes the genotype at variant *j* and *β_j_* is the corresponding effect size. The helper trait model extends the baseline model by incorporating *N_h_* helper traits as covariates: T2D ∼ Sex + Age + 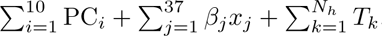. In total, we identified *N_h_* = 49 non-redundant helper traits. To characterize how predictive performance scales with the inclusion of helper traits, we fit a sequence of models in which the helper traits were added incrementally in their Pearson correlation with HbA1c rank order, *N_h_* = 1, 2*, . . .,* 49, enabling us to quantify the marginal contribution of additional, increasingly redundant predictors.

#### Model fitting and machine learning approaches

We performed predictive modeling on an independent dataset of UKB individuals consisting of 81,741 controls and 5,876 cases. This dataset was partitioned into three non-overlapping subsets: 20% for assessing general correlation structure among helper traits, 60% for training, and 20% for testing. To maximize predictive performance, we evaluated both linear and non-linear prediction models for T2D. Linear models treat genetic predictors and helper traits as additive covariates, while non-linear models, such as gradient boosting [73] and neural networks [74], capture higher-order interactions among predictors. We leveraged Automated Machine Learning (AutoML) to systematically explore predictive model architectures. Specifically, we used AutoGluon [41], a state-of-the-art automated machine learning framework that enables efficient exploration of complex model spaces and the identification of optimal predictive models. Unlike conventional approaches that rely on manually selecting a single model and tuning hyperparameters, AutoGluon systematically explores a wide range of machine learning algorithms, including XGBoost, LightGBM [75], deep neural networks, and ensemble methods. By automating hyperparameter tuning and model selection, Auto-Gluon ensures that our AUC-ROC scores reflect strong performance across multiple methodologies.

## Supplementary information

**Additional file 1**

File name: Additional file 1.pdf

File format: PDF (.pdf)

Title: Supplementary methods and figures

Description: Supplementary Methods 1, the redundancy-filtering algorithm used to select “tag helper traits”; Supplementary Methods 2, the derivation of the closed-form expression for the predictive gain; and Supplementary Figures S1–S4.

## Declarations

### Ethics approval and consent to participate

UK Biobank has ethical approval from the North West Multicentre Research Ethics Committee as a Research Tissue Bank (REC reference 21/NW/0157, renewing the original approval 11/NW/0382), under which the analyses reported here were conducted; no additional ethical approval was required. All UK Biobank participants provided written informed consent. This research was conducted using the UK Biobank Resource under application number 104628.

## Consent for publication

Not applicable.

## Availability of data and materials

Individual-level UK Biobank data are available to approved researchers through the UK Biobank; the analyses reported here were conducted under application number 104628. The theoretical and simulation code supporting the conclusions of this article, which reproduces Figures 2, 3, 4, S1, S2 and S3, is publicly available at https://github.com/AkeyLab/correlated-traits-prediction under the MIT licence.

Helper-trait predictive gain calculator (this study), https://akeylab.github.io/correlated-traits-prediction/

UK Biobank, https://www.ukbiobank.ac.uk

REGENIE, https://rgcgithub.github.io/regenie/

PLINK 2.0, https://www.coggenomics.org/plink/2.0/

AutoGluon, https://auto.gluon.ai/

## Authors’ contributions

KZ and JMA conceived and designed the study. KA and JMA developed the statistical framework, and KZ implemented the simulations and performed data analysis. KZ and JMA wrote the manuscript. RB contributed to the data curation and analysis. JMA supervised the study. All authors reviewed and approved the final manuscript.

## Acknowledgements.

We would like to thank the members of the Akey and Storey laboratories for helpful comments and feedback on this work.

