## Supplement File 1 for "A Principled Framework for Using Correlated Traits to Improve Risk Prediction"

### Additional file 1 — Supplementary methods and figures

**Abstract**

Supplementary material for "A Principled Framework for Using Correlated Traits to Improve Risk Prediction". Supplementary Methods 1 gives the redundancy-filtering algorithm used to select "tag helper traits". Supplementary Methods 2 derives the closed-form expression for the predictive gain in terms of heritabilities and genetic and environmental correlations. Supplementary Figures S1–S4 report the simulations validating that expression and the additional parameter regimes referred to in the main text.

#### Contents

**Supplementary Methods 1: Redundancy filtering algorithm** **2**

Supplementary Methods 2: Closed-form expression for the predictive gain 3

Supplementary Figures 4

### Supplementary Methods 1: Redundancy filtering algorithm

**Algorithm 1** Correlation-Based Feature Selection with Redundancy Filtering ("Tag Helper Traits")

**Input:**

$$\begin{aligned} T &= \{t_1, t_2, \ldots, t_n\}: \text{ candidate features, ranked by abs. corr. w/ HbA1c,} \\ \tau &= \text{pairwise correlation threshold } (\tau = 0.6 \text{ in this study}), \\ k &= \text{the number of selected features,} \\ C(i,j) &= \text{Pearson correlation between } t_i \text{ and } t_j \end{aligned}$$

**Output:**

$$S = \text{final set of selected ("tag") features}$$

1: $S \leftarrow \emptyset$
2: Add the top-ranked feature (highest absolute correlation to HbA1c) from $T$ to $S$
3: **for** each remaining feature $t_i$ in $T$ (in descending order of correlation) **do**
4: Compute $C(t_i, t_j)$ for every $t_j \in S$
5: **if** $|C(t_i, t_j)| < \tau$ for all $t_j \in S$ **then**
6: Add $t_i$ to $S$
7: **end if**
8: **if** $k$ is specified **and** $|S| = k$ **then**
9: **break**
10: **end if**
11: **end for**
12: **return** $S$

### Supplementary Methods 2: Closed-form expression for the predictive gain

For notational convenience, let $H_t = h_t^2$ and $H_h = h_h^2$ denote the target and helper heritabilities. For standardized phenotypes, the population phenotypic correlation between the target and helper traits is

$$\rho_p = \rho_g \sqrt{H_t H_h} + \rho_e \sqrt{(1 - H_t)(1 - H_h)}. \tag{1}$$

Because the baseline predictor $\boldsymbol{B}$ is constructed from the standardized target-trait genetic value and independent noise as in Equation (**??**) of the main text, its correlation with the target trait is

$$r_{tB} = \mathrm{Cor}(\boldsymbol{Y}_t, \boldsymbol{B}) = \sqrt{R^2_{\mathrm{base}}}, \tag{2}$$

and, using the same construction, its correlation with the helper trait is

$$r_{hB} = \mathrm{Cor}(\boldsymbol{B}, \boldsymbol{Y}_h) = \rho_g \sqrt{\frac{R^2_{\mathrm{base}} H_h}{H_t}}. \tag{3}$$

The incremental variance explained by adding the helper trait to the baseline predictor is

$$\Delta R^2 = \frac{(r_{th} - r_{tB}\, r_{hB})^2}{1 - r_{hB}^2}, \tag{4}$$

and therefore the relative predictive gain is

$$R^2_{\mathrm{gain}} = \frac{(r_{th} - r_{tB}\, r_{hB})^2}{R^2_{\mathrm{base}}\,(1 - r_{hB}^2)}. \tag{5}$$

At the population level, $r_{th}$ is replaced by $\rho_p$. Substituting the expressions for $r_{tB}$ and $r_{hB}$ gives

$$R^2_{\text{gain}} = \frac{\left[\rho_p - \rho_g\, R^2_{\text{base}} \sqrt{\dfrac{H_h}{H_t}}\right]^2}{R^2_{\text{base}} \left[1 - \rho_g^2\, R^2_{\text{base}}\, \dfrac{H_h}{H_t}\right]}. \tag{6}$$

Substituting the expression for $\rho_p$ yields a formula entirely in terms of the simulation inputs,

$$R^2_{\text{gain}} = \frac{\left[\rho_g \sqrt{\dfrac{H_h}{H_t}}\,\big(H_t - R^2_{\text{base}}\big) + \rho_e \sqrt{(1 - H_t)(1 - H_h)}\right]^2}{R^2_{\text{base}} \left[1 - \rho_g^2\, R^2_{\text{base}}\, \dfrac{H_h}{H_t}\right]}. \tag{7}$$

$$R^2_{\text{gain}} = \frac{\left[\rho_g \sqrt{\dfrac{h_h^2}{h_t^2}}\,\big(h_t^2 - R^2_{\text{base}}\big) + \rho_e \sqrt{(1 - h_t^2)(1 - h_h^2)}\right]^2}{R^2_{\text{base}} \left[1 - \rho_g^2\, R^2_{\text{base}}\, \dfrac{h_h^2}{h_t^2}\right]}. \tag{8}$$

Restoring the original notation ($H_t = h_t^2$, $H_h = h_h^2$), Equation (8) provides an equivalent closed-form expression for $R^2_{\text{gain}}$ as defined in Equation (**??**) of the main text, parameterized in terms of target- and helper-trait heritabilities, genetic and environmental correlations, and baseline predictive performance.

### Supplementary Figures

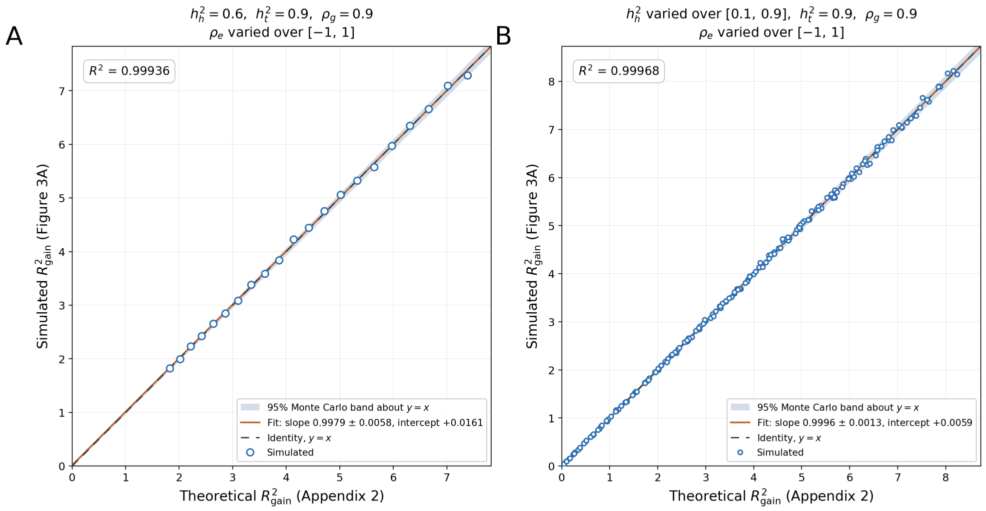


**Fig. S1 Simulations Validate the Closed-Form Expression for the Predictive Gain.** The relative gain in prediction accuracy, $R^2_{\text{gain}}$, evaluated in simulated data, is plotted against the closed-form expression derived in Supplementary Methods 2 below. Throughout, the target heritability is fixed at $h_t^2 = 0.9$, the genetic correlation at $\rho_g = 0.9$, and the baseline predictor captures a fraction $\alpha = 0.1$ of the target heritability, matching the design of Figure **??**A of the main text. (**A**) fixes the helper heritability ($h_h^2$) at 0.6 and varies the environmental correlation ($\rho_e$) over $[-1, 1]$, giving 21 points. (**B**) additionally varies the helper heritability ($h_h^2$) over $[0.1, 0.9]$, giving $21 \times 9 = 189$ points. Open circles represent the simulated data, each an average over 30 replicates, and the shaded bands indicate the 95% Monte Carlo agreement band about the identity line. Dashed black lines denote the identity $y = x$ and solid orange lines denote the least-squares fit, whose slope and intercept are given in the legend.

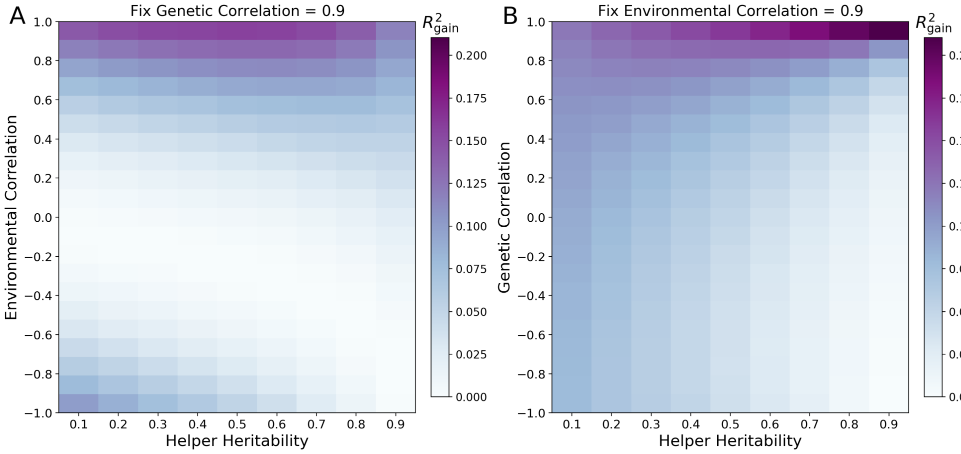


**Fig. S2 Gain in predictive accuracy for high-heritability target traits with strong baseline predictive performance.** (**A**) fixes the genetic correlation ($\rho_g$) at 0.9 and varies the environmental correlation ($\rho_e$) and helper heritability ($h_h^2$). (**B**) fixes the environmental correlation ($\rho_e$) at 0.9 while varying the genetic correlation ($\rho_g$) and helper heritability ($h_h^2$).

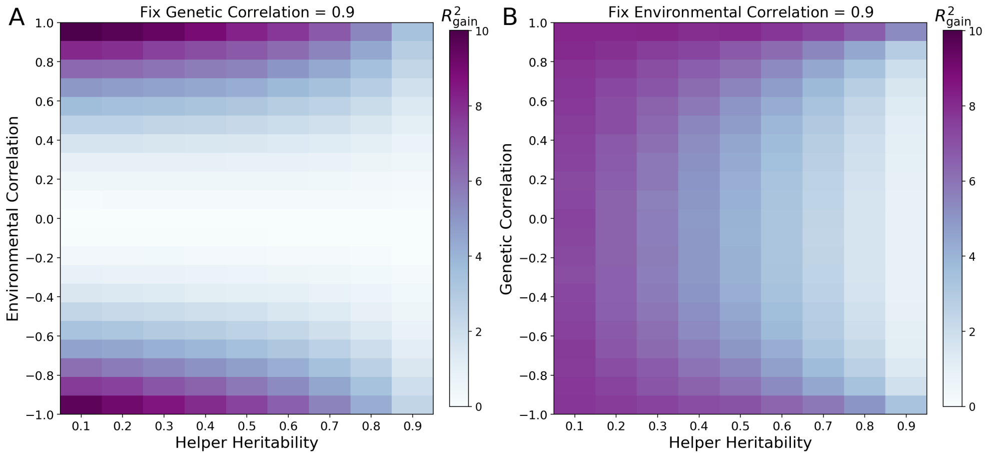


**Fig. S3 Gain in predictive accuracy for low-heritability target traits when $\alpha = 0.9$.** Here the baseline predictor captures most of the target heritability, but because $h_t^2 = 0.1$ it still explains only $R^2_{\text{base}} = 0.09$ of phenotypic variance. (**A**) fixes the genetic correlation ($\rho_g$) at 0.9 and varies the environmental correlation ($\rho_e$) and helper heritability ($h_h^2$). (**B**) fixes the environmental correlation ($\rho_e$) at 0.9 while varying the genetic correlation ($\rho_g$) and helper heritability ($h_h^2$).

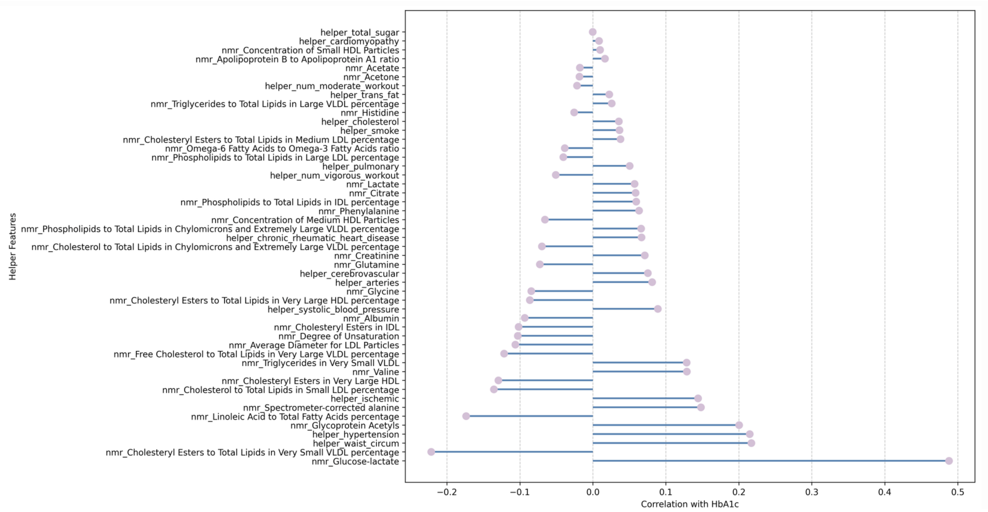


**Fig. S4 Correlation of "Tag Helper Traits" with HbA1c.** The 49 features shown here met the correlation threshold and redundancy criteria. Each bar corresponds to a single "tag" trait, with the length and direction indicating its correlation with HbA1c (negative on the left, positive on the right). Pink circles denote each trait's correlation point estimate. The list includes NMR-based lipid and metabolite measures as well as clinical and lifestyle-related helper traits, collectively illustrating diverse biological contributors to glycemic regulation and Type II Diabetes risk.
